# Exon capture optimization in large-genome amphibians

**DOI:** 10.1101/021253

**Authors:** Evan McCartney-Melstad, Genevieve G. Mount, H. Bradley Shaffer

## Abstract

**Background:** Gathering genomic-scale data efficiently is challenging for non-model species with large, complex genomes. Transcriptome sequencing is accessible for even large-genome organisms, and sequence capture probes can be designed from such mRNA sequences to enrich and sequence exonic regions. Maximizing enrichment efficiency is important to reduce sequencing costs, but, relatively little data exist for exon capture experiments in large-genome non-model organisms. Here, we conducted a replicated factorial experiment to explore the effects of several modifications to standard protocols that might increase sequence capture efficiency for large-genome amphibians.

**Methods:** We enriched 53 genomic libraries from salamanders for a custom set of 8,706 exons under differing conditions. Libraries were prepared using pools of DNA from 3 different salamanders with approximately 30 gigabase genomes: California tiger salamander (*Ambystoma californiense*), barred tiger salamander (*Ambystoma mavortium*), and an F1 hybrid between the two. We enriched libraries using different amounts of c_0_t-1 blocker, individual input DNA, and total reaction DNA. Enriched libraries were sequenced with 150 bp paired-end reads on an Illumina HiSeq 2500, and the efficiency of target enrichment was quantified using unique read mapping rates and average depth across targets. The different enrichment treatments were evaluated to determine if c_0_t-1 and input DNA significantly impact enrichment efficiency in large-genome amphibians.

**Results:** Increasing the amounts of c_0_t-1 and individual input DNA both reduce the rates of PCR duplication. This reduction led to an increase in the percentage of unique reads mapping to target sequences, essentially doubling overall efficiency of the target capture from 10.4% to nearly 19.9%. We also found that post-enrichment DNA concentrations and qPCR enrichment verification were useful for predicting the success of enrichment.

**Conclusions:** Increasing the amount of individual sample input DNA and the amount of c_0_t-1 blocker both increased the efficiency of target capture in large-genome salamanders. By reducing PCR duplication rates, the number of unique reads mapping to targets increased, making target capture experiments more efficient and affordable. Our results indicate that target capture protocols can be modified to efficiently screen large-genome vertebrate taxa including amphibians.

## Background

Reduced representation sequencing technologies enrich DNA libraries for selected genomic regions, allowing researchers to attain higher sequencing depth over a predetermined subset of the genome for a given cost. Several techniques are now in widespread use in population genetics and evolutionary biology. The most popular of these include RAD-tag sequencing (which targets anonymous loci flanking restriction enzyme sites) [1] and target-enrichment approaches such as ultra-conserved element (UCE) sequencing (which targets regions of the genome that are highly-conserved between species) [2] and exome/exon sequencing (which target genomic regions that are expressed as RNAs).

These methods are all extremely useful for different purposes. RAD-tag sequencing is a cost-effective strategy for collecting information on thousands of anonymous loci for individuals within a population, but suffers from bias and large amounts of missing data, especially when divergent individuals are analyzed [3]. UCE sequencing, conversely, is designed to generate relatively complete datasets across distantly related species using a single test panel [2], but the biological function of these conserved loci are mostly unknown.

Exon capture differs from RAD-tag sequencing in that it targets predetermined sequence regions, and is distinct from UCE sequencing in that it targets known gene regions that are often assumed to be functionally important. As such, exon capture is a promising technology for gathering large amounts of targeted genomic data for population-level studies exploring patterns of population structure and natural selection [4–6]. It is particularly useful for collecting data from species without assembled reference genomes, as the prerequisite genomic information may be gathered from existing collections of expressed sequence tag (EST) sequences or transcriptome sequencing [7, 8]. Enrichment of exon sequences has been performed in multiple non-model species, with applications ranging from investigating genotype/phenotype associations to population genetics and phylogenetic inference [2, 7–9].

The molecular laboratory principles of UCE sequencing and exon capture sequencing are the same. Both procedures rely on the hybridization of synthetic biotinylated RNA or DNA probes to library fragments from samples of interest. After hybridization, the biotin on these probes is bound to streptavidin molecules attached to magnetic beads, allowing the target sequences to be magnetically captured, and all non-hybridized DNA is washed away. Unfortunately, capture of off-target DNA can happen for several reasons, and can drastically reduce the efficiency of sequencing [10]. Because library fragments are often longer than the probe sequences, part of the hybridized library fragment is usually free to bind to other molecules in the pool. Since repetitive DNA sequences are by definition present at high concentrations in large-genome organisms, if this exposed region is from a repetitive element it has a high probability of binding to another such fragment and pulling it through to the final library pool. Adapter sequences are also present at very high concentrations, presenting another opportunity for molecules to bind to captured fragments, creating “daisy chains” of random library molecules. To mitigate these factors, several “blockers,” designed to hybridize to these regions before the biotinylated probes are able to, are typically added to target capture reactions. One such blocker, c_0_t-1, is a solution of high-copy repetitive DNA fragments that hybridizes with repetitive library fragments and blocks them from attaching to captured fragments. For large-genome amphibians, repetitive elements are present at an even higher concentration than normal [11], and we hypothesize that increasing the amount of c_0_t-1 in solution may improve hybridization efficiency. This process is shown in Figure 1.

**Figure 1.**
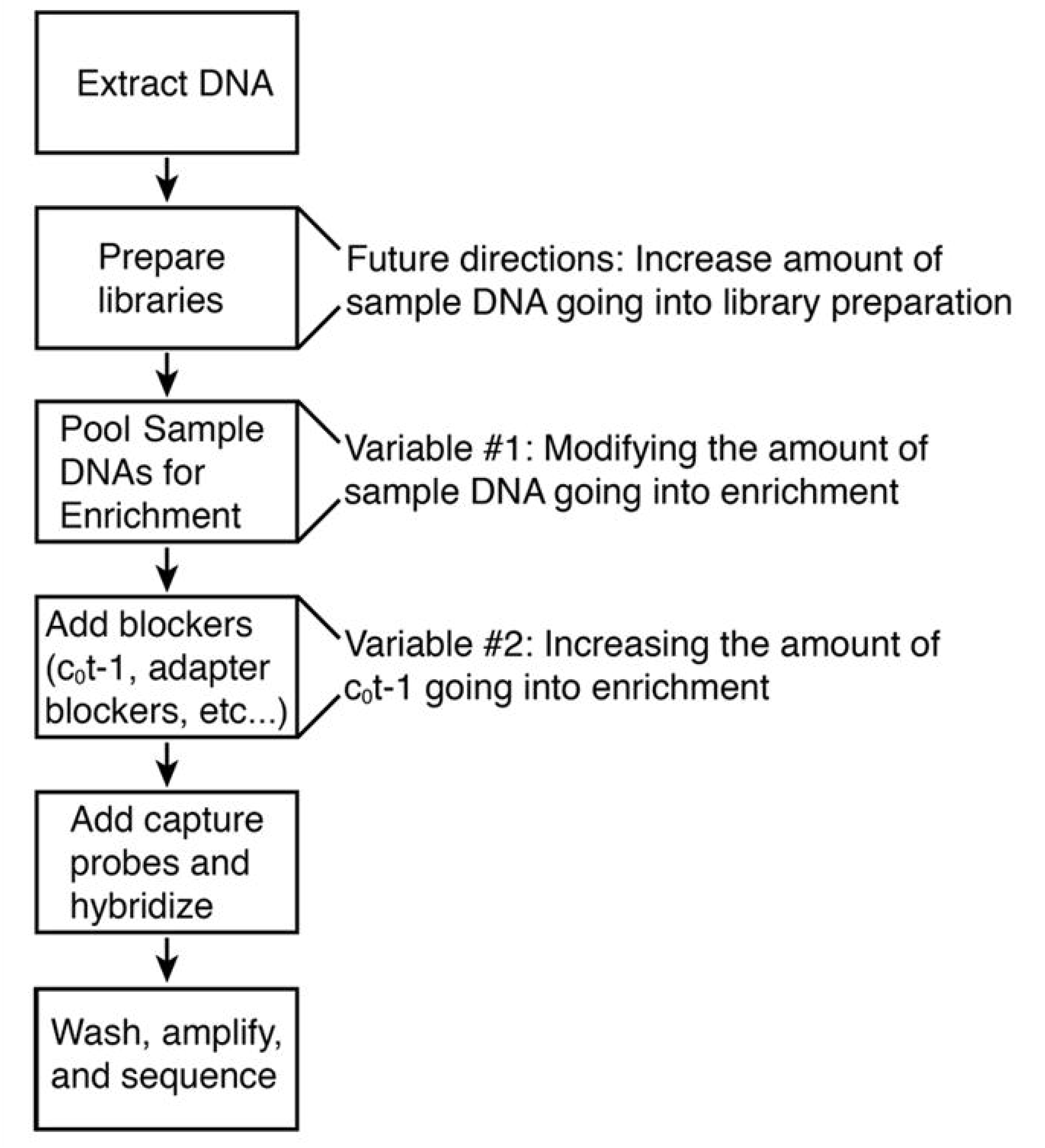
Flow chart depicting target enrichment process and key steps affected by experimental variables.

Relatively few exon capture studies have been performed in amphibians [but see 9], likely reflecting the reticence of many biologists to apply genomic approaches to their large, highly repetitive genomes. While these large genomes, ranging up to 117 gigabases [12], currently render full-genome sequencing approaches untenable, exon capture is well-suited to bridge the gap between single-locus comparative studies and whole-genome analyses for these and other large-genome diploid species. Several amphibian species have large collections of EST sequences available [13–15], and sequencing of cDNA libraries with *de novo* transcriptome assembly is becoming increasingly accessible for species that currently lack such resources.

Laboratory costs of exon capture experiments hinge largely on the efficiency of the enrichment process. Increasing the percentage of reads “on target” (sequence reads that align to regions targeted in the capture array) directly reduces the amount of sequencing required to attain a desired coverage level. Off-target reads may be present for several reasons, including non-specific hybridization of capture probes to off-target regions, hybridization of off-target DNA to the ends of captured target fragments, and failure to wash away all DNA not hybridized to capture probes following enrichment [10]. This process may be particularly problematic in amphibians because their large genome size is often due to a massive increase in the amount of repetitive DNA [11], which leads to an greatly increased concentration of off target DNA in solution relative to on-target fragments.

We conducted a series of experiments that seek to optimize existing protocols for exon capture experiments for large-genome amphibians (and other taxa). Our focus is on three different *Ambystoma* salamanders—the California tiger salamander (*Ambystoma californiense*), the barred tiger salamander (*Ambystoma mavortium*), and an F1 hybrid between the two (*Ambystoma californiense x mavortium*, referred to as F1). Given the enormous size of their genomes (estimated at about 32 gigabases) and the observation that they, like many amphibians, have genomes that are rich in repetitive DNA, we altered the amount of c_0_t-1 blocker, under the assumption that highly-repetitive genomes may benefit from an increased amount of repetitive sequence blocker. We also manipulated the amount of individual input and total DNA in sequence capture reactions to manipulate the total number of copies of the genome, estimating tradeoffs among multiplexibility and enrichment efficiency to maximize the number of individuals that can be sequenced for each sequence capture reaction.

## Methods

### Array design and laboratory methods

We designed an array of 8,706 putative exons (8,706 distinct genes) using EST sequences from the closely-related Mexican axolotl (*Ambystoma mexicanum*) [17]. Mitochondrial sequence divergence between the California tiger salamander and the Mexican axolotl is approximately 6.4%, and is approximately 6.8% between the barred tiger salamander and Mexican axolotl [18], suggesting that less-diverged nuclear exons from the axolotl should serve as appropriate targets for our species. In our design, we attempted to avoid targeting regions that span exon/intron boundaries, as these targets have been found to be much less efficient [8]. Exon boundaries can be found by mapping EST sequences to a reference genome while allowing for long gaps that represent introns. However, no salamander genome is currently available, and the two available frog genomes (*Xenopus tropicalus* [19] and *Nanorana parkeri* [20]) last shared a common ancestor with salamanders approximately 290 million years ago [21]. To account for this, we developed a comparative method for conservatively predicting intron splice sites within EST sequences (unpublished data). Target sequences were an average of 290 bp in length (minimum length=88 bp, maximum=450 bp, standard deviation=71 bp), for a total target region length of 2.53 megabases. A total of 39,984 100bp probe sequences were tiled across these target regions at an average of 1.8X tiling density. These probes were synthesized as biotinylated RNA oligos in a MYbaits kit (MYcroarray, Ann Arbor, MI).

We extracted genomic DNA from three individual salamanders--one California tiger salamander (*Ambystoma californiense #*HBS127160—CTS), one barred tiger salamander (*Ambystoma mavortium* #HBS127161—BTS), and one F1 hybrid between the two species (#HBS109668)—using a salt extraction protocol [22] and several independent extractions of each individual to attain the amount needed for preparing several libraries. Extractions were then combined into pools to draw from for library preparations. Two of these pools consisted of pure California tiger salamander DNA or pure F1 DNA and are labeled CTS and F1, respectively. The third pool, which was intended to be pure BTS, was found to consist of roughly 70% barred tiger salamander DNA and 30% California tiger salamander DNA, apparently due to a pooling error (later verified through re-extraction of the original tissues and Sanger sequencing). We refer to this pool as BTS*, and treat it as a third sample in our experimental design. DNA was diluted to 20 ng/μL and sheared to roughly 500bp on a BioRupter (Diagenode, Denville, NJ). For each of the 53 individual library preparations (Table 1), we used roughly 450 ng of DNA for library preparations. Standard Illumina library preparations (end repair, A-tailing, and adapter ligation) were performed using Kapa LTP library preparation kits (Kapa Biosystems, Wilmington, MA). Samples were dual-indexed with 8 bp indices that were added via PCR (adapters from Travis Glenn, University of Georgia). Following library preparation we performed a double-sided size selection with SPRI beads [23] to attain a fragment size distribution centered around 400 bp and ranging from 200bp to 1,000 bp. Species-specific c_0_t-1 was prepared using DNA extracted from a California tiger salamander and a single-strand nuclease as follows: First, extracted DNA was treated with RNase and brought to 500 μL at 1,000 ng/μL in 1.2X SSC. This DNA was then sheared on a BioRuptor (Diagenode, Denville, NJ) to roughly 300bp. Next, the solution was denatured at 95C for 10 minutes, then partially renatured at 60C for 5 minutes and 45 seconds, placed on ice for two minutes, then put in a 42C incubator. A preheated 250 μL aliquot of S1 nuclease (in buffer) was then added to the partially-renatured DNA and incubated for 1 hour at 42C. The DNA was then precipitated with 75 μL of 3M sodium acetate and 750 μL isopropanol and centrifuged for 20 minutes at 14,000 RPM at 4C. Isopropanol was then removed and the pellet was washed with 500 μL cold 70% ethanol, centrifuged again at 14,000 RPM for 10 minutes (4C), and dried following ethanol removal. We rehydrated this pellet with 50 μL of 10 mM Tris-HCl, pH 8, and dried down to the appropriate concentration (for 1X c_0_t-1, 500 ng/μL; for 6X and 12X c_0_t-1 1,000 ng/μL).

**Table 1.**
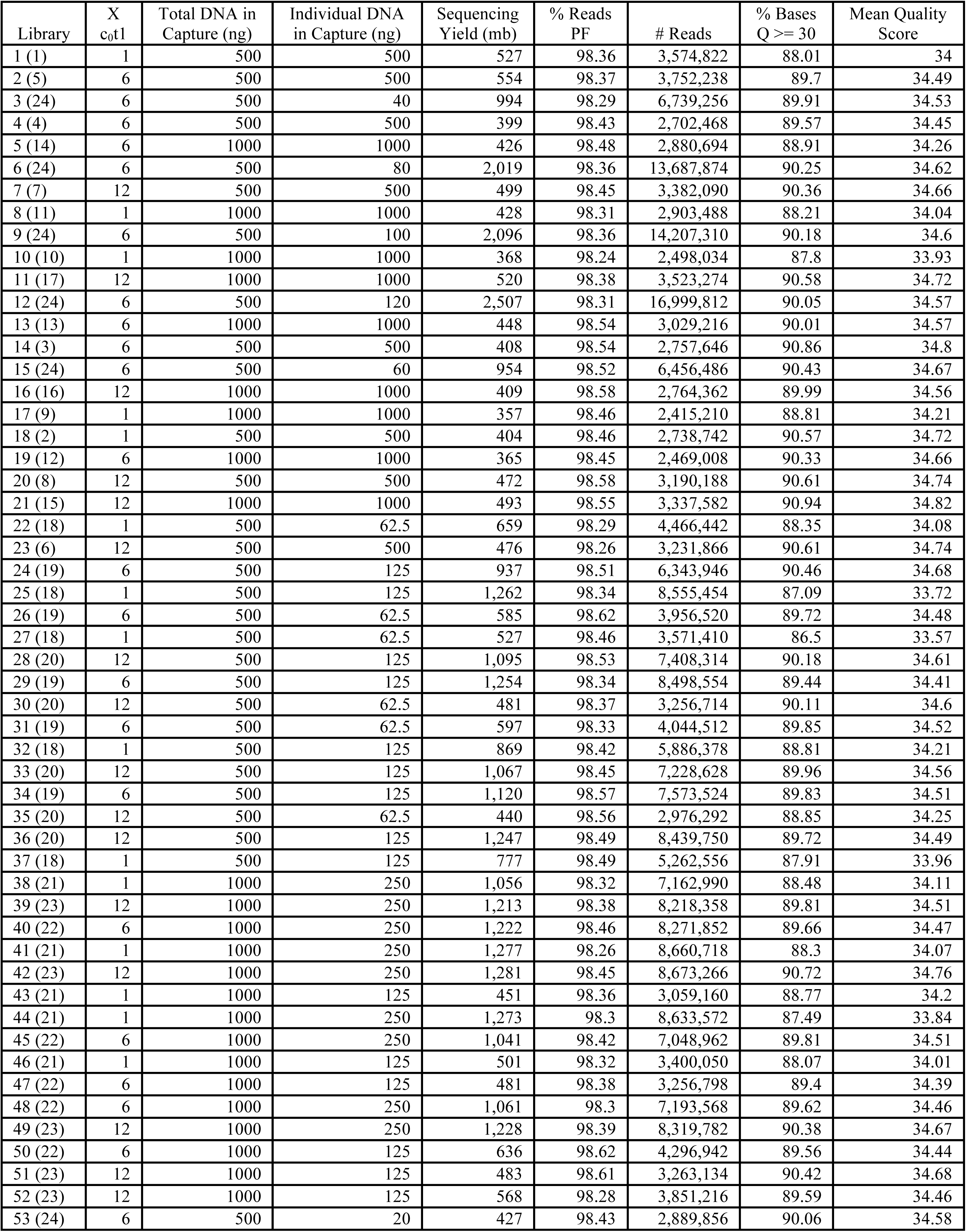
Individual libraries (1-53), their treatment levels, and description of yields and sequencing statisttics. Number in parenthesis in library (first column) is the enrichment (1-24), and shows how libraries were pooled. For example, 22(18) and 25(18) indicates that libraries 22 and 25 were pooled into a single tube (number 18) prior to enrichment.

We then multiplexed prepared libraries into capture reactions (Table 1). Total DNA input into the sequence capture was either 500 ng or 1,000 ng, and individual library input DNA for multiplexing ranged from 20 to 1,000 ng (Table 1). The repetitive DNA blocker c_0_t-1 was added to the 24 different capture reactions in one of three amounts—2,500 ng, 15,000 ng, or 30,000 ng, corresponding to 1X, 6X, and 12X protocol recommendation. Libraries were enriched using the MYbaits protocol (version 2.3.1), hybridizing probes for 24.5 hours and implementing the optional high-stringency washes. Following the three wash steps in the MYbaits protocol, we amplified the remaining enriched DNA (with streptavidin beads still in solution) using 14 cycles of PCR. Multiple separate PCR reactions were performed for each capture reaction, which were subsequently pooled after amplification to reduce PCR amplification bias [24].

Post-capture, post-PCR libraries were quantitated and characterized with qPCR using the Kapa Illumina library quantification kit (PicoGreen® Life Technologies, Grand Island, NY and Kapa Biosystems, Wilmington, MA) on a LightCycler 480 (Roche, Basel, Switzerland). We also visualized fragment size distributions using a BioAnalyzer 2100 DNA HS chip (Agilent, Santa Clara, CA). All capture reactions were tested for preliminary evidence of enrichment via qPCR. We developed five primer pairs derived from different test loci chosen from our targets as positive controls, and one primer pair derived from a mitochondrial locus we were not targeting as a negative control. We used these to measure the relative concentrations of target molecules in solution by calculating the mean number of cycles required for qPCR reactions to reach the crossing point (C_p_) in libraries pre and post enrichment. Changes in (C_p_) were measured for each test locus for all samples and averaged across all five test loci. For targeted loci, we expected that the number of cycles needed to reach this point would decrease, because target sequences would be present in higher concentrations. Conversely, we expected the number of cycles for the mitochondrial DNA locus to increase after enrichment, because that sequence was not targeted and we expected its concentration to decrease.

All capture reactions were then combined together for sequencing on an Illumina HiSeq 2500 with 150bp paired-end reads. Reactions were pooled such that all individual libraries would receive at least 1.5 million reads. Because some capture reactions contained samples with more DNA compared to other samples in the pool, some capture reactions were assigned more of the sequencing lane than others (Table 1). Sample pooling and sequencing was performed at the Vincent J. Coates Genomics Sequencing Laboratory at UC Berkeley.

### Genetic data analysis

Demultiplexed reads were checked for adapter contamination and quality trimmed using Trimmomatic 0.32 [25]. Quality trimming was performed using several criteria. First, leading base pairs with a phred score less than 5 were removed. Next, trailing (3’) base pairs with a phred score less than 15 were removed. Finally, we used a four base pair sliding window (5’ to 3’), trimming all trailing bases when the average phred score within that window dropped below 20. We discarded all reads under 40 bp after trimming, and overlapping reads were merged using fastq-join [26].

Genetic data from all of the Califonia tiger salamander libraries were combined for assembly to create the most complete possible single-species *de novo* assembly of our target regions. Targets were *de novo* assembled using the Assembly by Reduced Complexity (ARC) pipeline [27]. This assembly pipeline separates reads that align to target regions and performs small, target-specific *de novo* assemblies on these read pools. Each assembled contig then replaces its original target sequence, and the process is repeated iteratively. Within ARC, read mapping was performed using bowtie2 [28], error correction with BayesHammer [29], and assemblies were generated using SPAdes [30]. The ARC pipeline was run for six iterations, which was enough to exhaust all of the reads assignable to most targets.

Following assembly, all contigs were compared against the original target sequences using blastn [31], and reciprocal best blast hits (RBBHs) were found [32]. Chimeric assemblies are pervasive and problematic for studies that involve *de novo* assembly of target sequences, because they can insert repetitive sequences into the contigs, making it appear that many reads are mapping to a target when those reads are actually from repetitive regions in the genome (for instance, see the coverage across the non chimera-masked contig in Figure 2). To attempt to reduce the presence of chimeric assemblies and repetitive sequences in our data, the RBBHs were blasted to themselves (blastn e-value of 1e-20), and base pairs in sequence regions that positively matched other targets were replaced with N’s. These chimera-masked RBBHs served as our final assembled target set.

**Figure 2.**
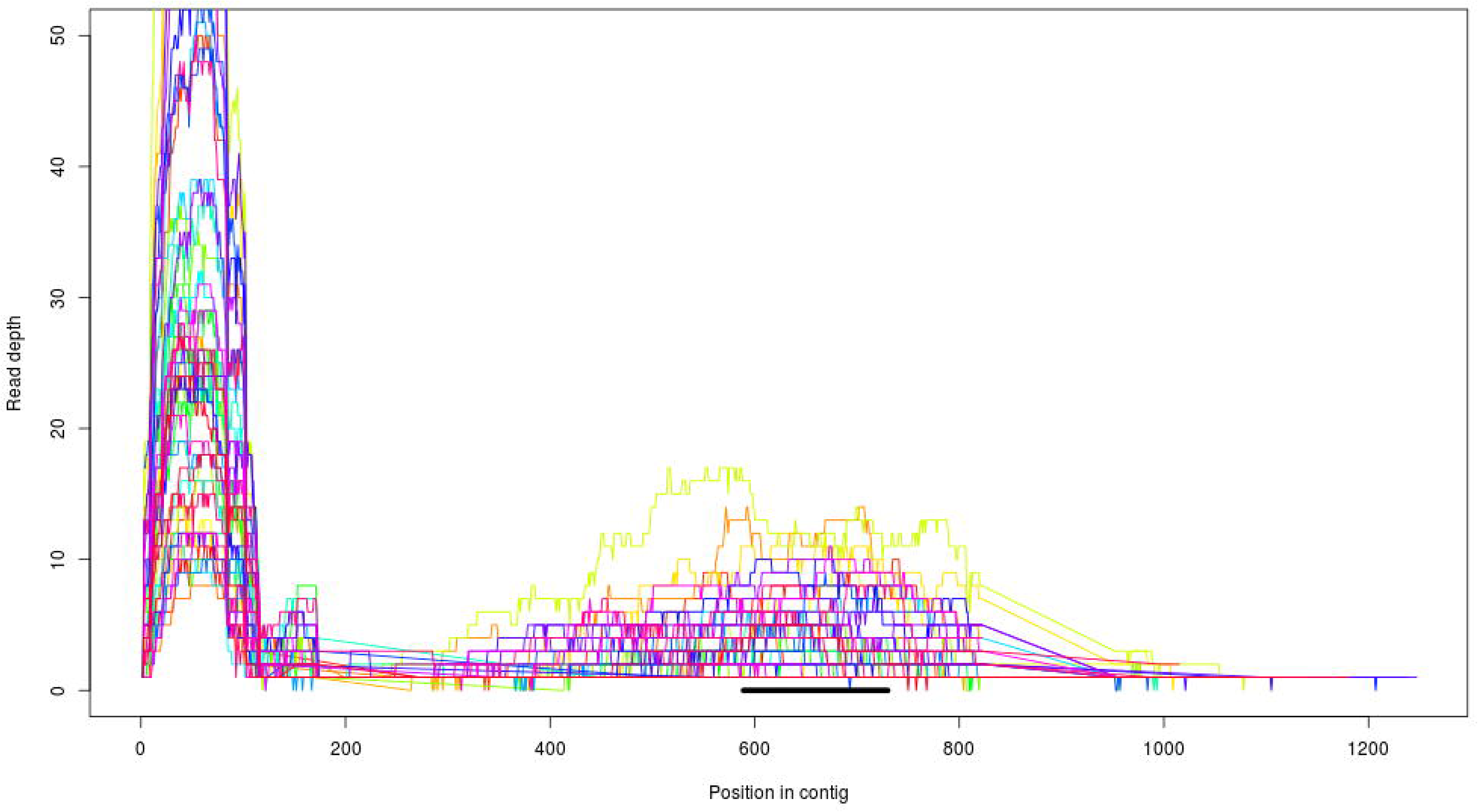
Coverage across a sample target. The black bar on the bottom corresponds to the target region from which probes were synthesized. Each line represents a single library, and each library is shown in a different color. There are two peaks of coverage, one centered on the target region, and a much higher spike of coverage at the left edge of the contig, likely corresponding to a repetitive region in the genome. The latter type of spikes are reduced through the chimera-filtering steps described in the text.

After assembly, reads from each individual were mapped against the chimera-masked RBBH target set using bwa mem [33]. BAM file conversion, sorting, and merging was done using SAMtools v1.0 [34]. PCR duplicates were marked using picard tools v. 1.119 (http://broadinstittute.github.io/picard) which finds reads or read pairs that have identical 5’ and 3’ mapping coordinates, with the reasoning that two chromosome copies are unlikely to shear in the exact same positions during random sonication. Under this assumption, reads or read pairs that have identical 5’ and 3’ mapping coordinates likely result from sequencing multiple amplified copies of the same original DNA molecule, which is undesirable. Finally, mapping rates and PCR duplication rates were inferred by counting the relevant SAM flags using SAMtools flagstat [34].

In addition to measuring the total percentage of unique reads that mapped to target regions, target-level performance was also evaluated. Because most targets showed a characteristic peak of read depth centered over the middle of the target where probes were tiled, and because a few targets maintained confounding repetitive sequences at the periphery of the assembled contigs, we characterized the read depths of targets over bases that had direct overlap with our target probes. That is, for target-level metrics, we did not consider read depth for the flanking regions that are naturally appended to the ends of each target during the assembly process. For each individual library preparation, we calculated the average unique-read sequencing depth across a) the entire target regions and b) across the 100 bp window within each target that had the highest average coverage. For all read depth comparisons, depths were corrected for the total number of reads a library received in sequencing by multiplying by a scaling factor n_f_/n_i_, where n_f_ is the fewest number of reads received by any individual in the experiment and n_i_ is the number of reads received by the individual under consideration. Assembled target sequences less than 100 bp were not included in read depth calculations because 100 bp is significantly less than the average read length and these targets tended to recruit very few reads.

### Assessing the importantance of c_0_t-1 and individual input DNA amounts

Linear regression was used to quantify the relationships between c_0_t-1 and individual input DNA to the percentage of unique reads that mapped to targets. Because three different biological individuals were used for library preparations in this experiment, we also included the identity of the individual as a possible source of variation to explain enrichment efficiency. Models were built that included different combinations of c_0_t-1, individual input DNA, and the identity of the individual (CTS, BTS*, or F1) as predictor variables, and unique reads mapping to targets as the response variable. A similar approach was used to model the average sequencing depths across all targets. All models were evaluated by examining the regression coefficients, adjusted R^2^, and AIC values.

## Results

### Pre-sequencing library quantitation

DNA concentration yields for post-enrichment, post-PCR samples were lower than anticipated. After 14 PCR cycles, amplified enrichment pools contained an average of 279.5 ng of DNA (after amplifying 15 μL out of a total 33 μL in the post-enrichment pools with a 50 μL PCR reaction). One capture reaction (Library # 18, see Table 2) had a much higher yield after post-enrichment PCR (2,150 ng). Mean C_p_ in qPCR enrichment verification reactions decreased by an average of 9.1 cycles across the five test loci after enrichment, while the number of cycles required for amplification of a non-targeted negative control locus increased by an average of 2.17 cycles. We found a positive correlation between the mean change in C_p_ averaged across the five test loci and the raw percentage of reads on target after sequencing for each library (Figure 3, adjusted R^2^ = 0.1136, p = 0.00784), although the relationship was stronger between post-enrichment, post-PCR DNA concentration and raw mapping rate (Figure 4, adjusted R^2^ = 0.224, p = 0.000204).

**Figure 3.**
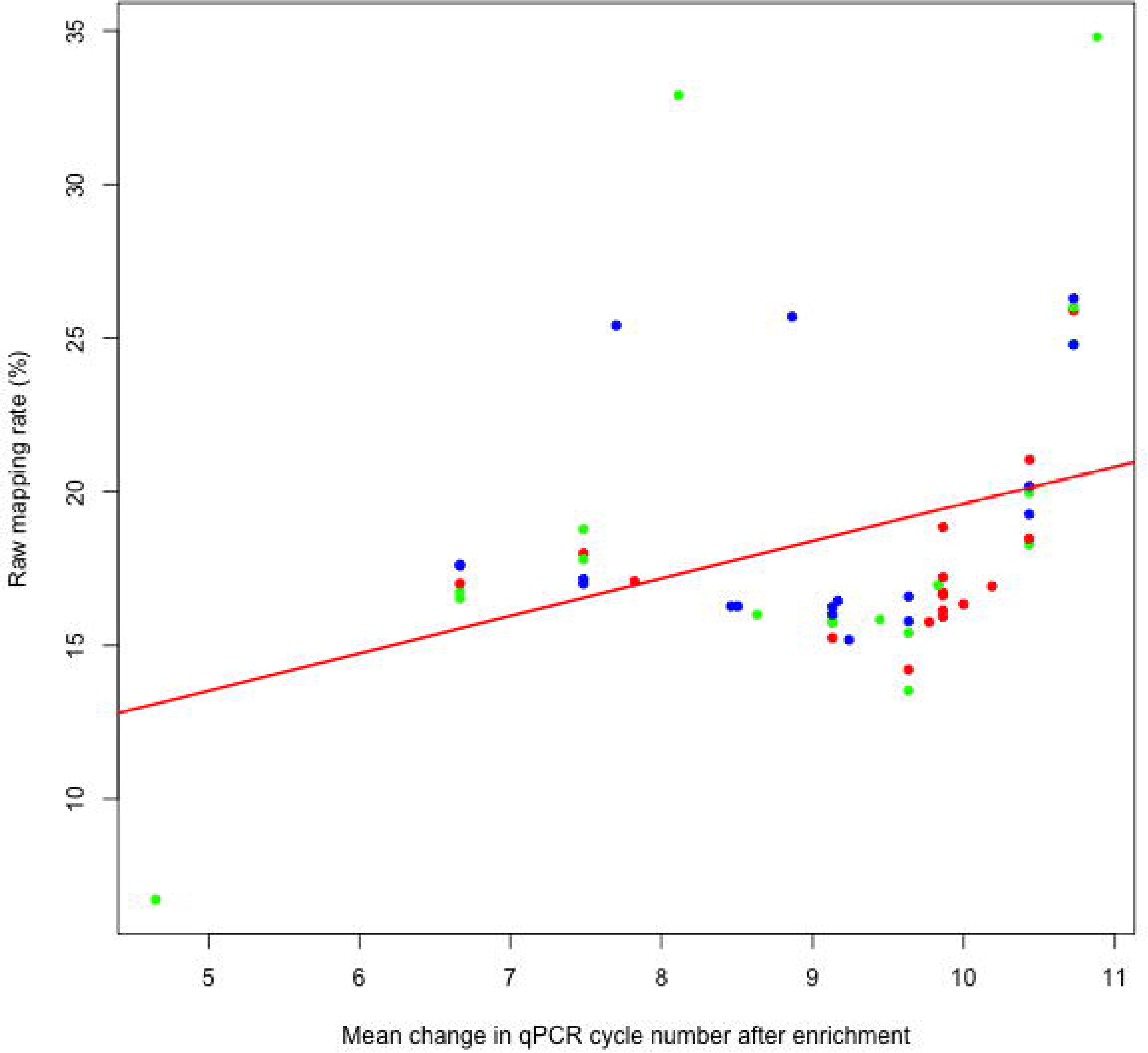
The change in raw mapping rate as a function of post-enrichment qPCR cycle number. Each dot is an individual library: blue=CTS, green=F1, red=BTS*. Adjusted R^2^ = 0.1136, p = 0.00784.

**Figure 4.**
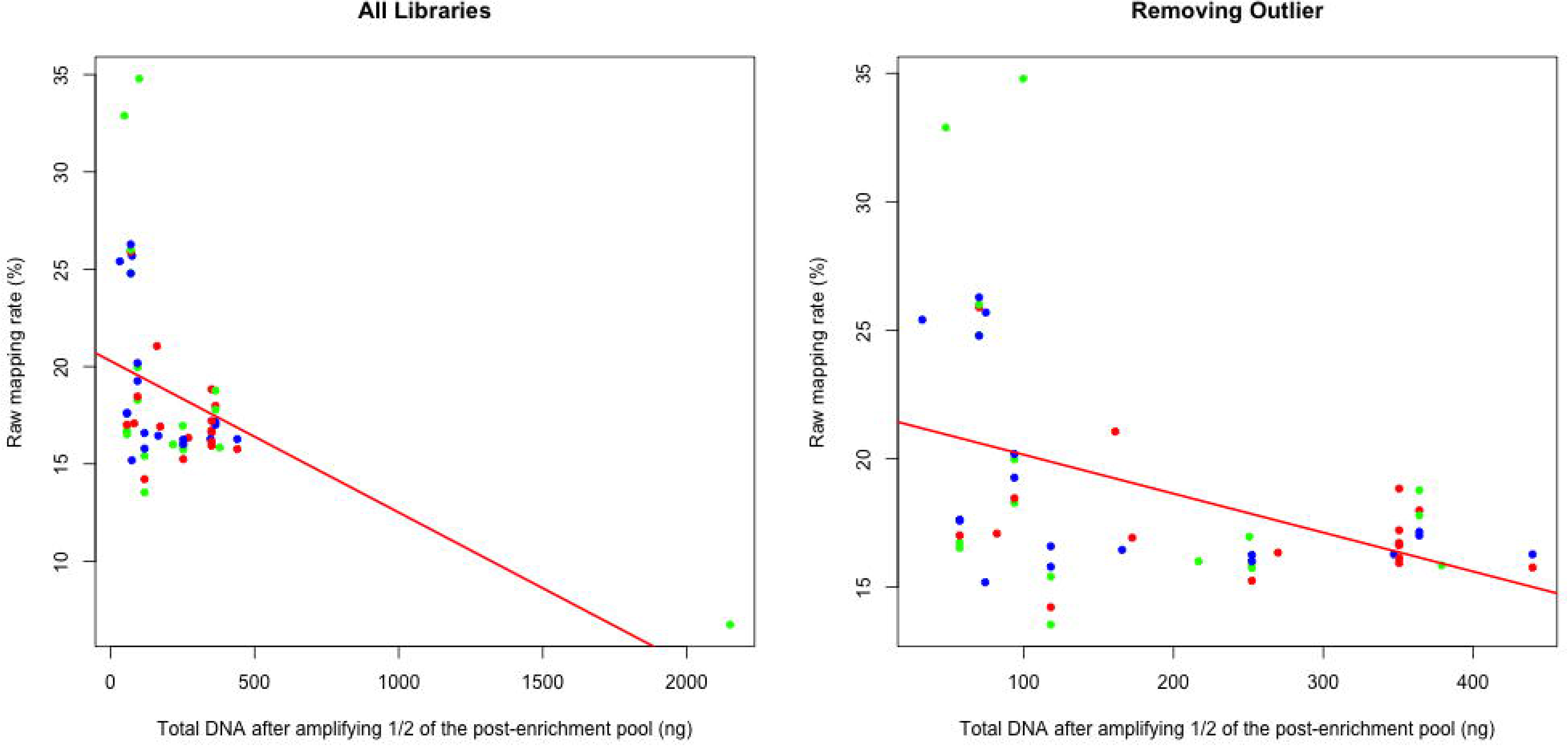
Relationship between post-enrichment DNA concentration and percentage of raw reads mapping to targets. Each dot is an individual library: blue=CTS, green=F1, red=BTS*. For the full dataset, adjusted R^2^ = 0.224, p = 0.000204. After removing the single F1 outlier, adjusted R^2^ =0 1732, p = 0.00126.

**Table 2.** Post-enrichment concentrations and sequencing efficiency results. Number in parenthesis in library name as in Table 1.

| Library # | Amount of DNA after post-enrichment PCR | Average change in qPCR cycle # | PCR duplication rate | Raw mapping rate | Unique read mapping rate | Average depth across target | Highest 100bp window ave. depth |
| --- | --- | --- | --- | --- | --- | --- | --- |
| 1 (1) | 74.1746508 | 9.2388 | 22.24% | 15.19% | 11.81% | 6.27 | 7.68 |
| 2 (5) | 47.82866 | 8.113 | 16.70% | 32.89% | 27.40% | 18.63 | 21.76 |
| 3 (24) | 350.64 | 9.864 | 30.26% | 16.72% | 11.66% | 6.58 | 7.74 |
| 4 (4) | 32.15888 | 7.6988 | 16.21% | 25.41% | 21.29% | 13.22 | 15.85 |
| 5 (14) | 99.542055 | 10.883 | 11.39% | 34.79% | 30.83% | 20.89 | 24.45 |
| 6 (24) | 350.64 | 9.864 | 33.19% | 16.63% | 11.11% | 6.32 | 7.31 |
| 7 (7) | 347.135 | 8.505 | 10.58% | 16.28% | 14.55% | 7.70 | 9.27 |
| 8 (11) | 216.706105 | 8.632 | 15.23% | 16.00% | 13.56% | 7.74 | 9.42 |
| 9 (24) | 350.64 | 9.864 | 33.57% | 15.93% | 10.59% | 5.94 | 6.86 |
| 10 (10) | 165.65566 | 9.1638 | 16.33% | 16.45% | 13.76% | 7.54 | 9.31 |
| 11 (17) | 379.095 | 9.446 | 9.21% | 15.84% | 14.38% | 7.78 | 9.30 |
| 12 (24) | 350.64 | 9.864 | 33.72% | 16.14% | 10.69% | 5.99 | 6.88 |
| 13 (13) | 74.65109 | 8.863 | 11.91% | 25.69% | 22.63% | 13.97 | 16.59 |
| 14 (3) | 81.971965 | 7.82 | 11.59% | 17.08% | 15.10% | 8.73 | 10.62 |
| 15 (24) | 350.64 | 9.864 | 29.10% | 18.83% | 13.35% | 7.86 | 9.20 |
| 16 (16) | 439.73 | 8.461 | 9.11% | 16.27% | 14.79% | 7.68 | 9.32 |
| 17 (9) | 172.37966 | 10.187 | 14.21% | 16.92% | 14.52% | 8.65 | 10.60 |
| 18 (2) | 2150.07215 | 4.652 | 19.62% | 6.74% | 5.42% | 0.27 | 0.47 |
| 19 (12) | 161.068535 | 10.435 | 8.73% | 21.05% | 19.22% | 11.91 | 14.30 |
| 20 (8) | 250.64 | 9.833 | 8.84% | 16.96% | 15.46% | 8.52 | 10.19 |
| 21 (15) | 269.695 | 9.999 | 8.51% | 16.34% | 14.95% | 8.43 | 10.09 |
| 22 (18) | 57.148215 | 6.668 | 47.05% | 16.72% | 8.86% | 4.69 | 5.72 |
| 23 (6) | 439.78 | 9.773 | 8.75% | 15.76% | 14.38% | 8.16 | 9.80 |
| 24 (19) | 117.984425 | 9.637 | 31.86% | 13.54% | 9.22% | 4.45 | 5.36 |
| 25 (18) | 57.148215 | 6.668 | 48.59% | 17.62% | 9.06% | 4.74 | 5.64 |
| 26 (19) | 117.984425 | 9.637 | 30.90% | 15.41% | 10.65% | 5.46 | 6.63 |
| 27 (18) | 57.148215 | 6.668 | 47.69% | 17.59% | 9.20% | 4.66 | 5.76 |
| 28 (20) | 93.68 | 10.432 | 19.83% | 18.28% | 14.65% | 8.37 | 9.72 |
| 29 (19) | 117.984425 | 9.637 | 33.69% | 16.59% | 11.00% | 5.67 | 6.70 |
| 30 (20) | 93.68 | 10.432 | 17.64% | 19.97% | 16.44% | 9.67 | 11.49 |
| 31 (19) | 117.984425 | 9.637 | 31.14% | 15.79% | 10.87% | 5.39 | 6.55 |
| 32 (18) | 57.148215 | 6.668 | 46.67% | 17.01% | 9.07% | 5.08 | 6.15 |
| 33 (20) | 93.68 | 10.432 | 21.03% | 19.26% | 15.21% | 8.55 | 9.96 |
| 34 (19) | 117.984425 | 9.637 | 32.57% | 14.22% | 9.59% | 4.96 | 5.90 |
| 35 (20) | 93.68 | 10.432 | 16.78% | 20.18% | 16.79% | 9.60 | 11.49 |
| 36 (20) | 93.68 | 10.432 | 20.87% | 18.46% | 14.61% | 8.67 | 10.05 |
| 37 (18) | 57.148215 | 6.668 | 46.14% | 16.53% | 8.90% | 4.82 | 5.83 |
| 38 (21) | 252.28847 | 9.128 | 28.10% | 15.24% | 10.96% | 6.09 | 7.24 |
| 39 (23) | 364.13 | 7.482 | 17.49% | 17.00% | 14.03% | 7.64 | 8.91 |
| 40 (22) | 70.102805 | 10.725 | 38.12% | 25.89% | 16.02% | 10.11 | 11.74 |
| 41 (21) | 252.28847 | 9.128 | 29.75% | 15.74% | 11.05% | 6.03 | 7.10 |
| 42 (23) | 364.13 | 7.482 | 16.69% | 17.99% | 14.98% | 8.82 | 10.22 |
| 43 (21) | 252.28847 | 9.128 | 25.29% | 15.92% | 11.89% | 6.36 | 7.78 |
| 44 (21) | 252.28847 | 9.128 | 30.89% | 16.01% | 11.06% | 5.75 | 6.81 |
| 45 (22) | 70.102805 | 10.725 | 35.17% | 26.01% | 16.86% | 10.52 | 12.25 |
| 46 (21) | 252.28847 | 9.128 | 27.01% | 16.25% | 11.86% | 6.15 | 7.53 |
| 47 (22) | 70.102805 | 10.725 | 34.28% | 24.80% | 16.30% | 10.11 | 12.06 |
| 48 (22) | 70.102805 | 10.725 | 37.39% | 24.78% | 15.52% | 9.33 | 10.92 |
| 49 (23) | 364.13 | 7.482 | 16.22% | 17.79% | 14.90% | 8.48 | 9.81 |
| 50 (22) | 70.102805 | 10.725 | 36.08% | 26.28% | 16.80% | 10.22 | 12.10 |
| 51 (23) | 364.13 | 7.482 | 11.98% | 18.77% | 16.52% | 9.45 | 11.23 |
| 52 (23) | 364.13 | 7.482 | 14.06% | 17.15% | 14.74% | 7.83 | 9.35 |
| 53 (24) | 350.64 | 9.864 | 26.20% | 17.21% | 12.70% | 6.98 | 8.46 |

### Sequence data

We generated 45,641,469,300 base pairs of sequence data in the form of 150bp paired-end reads. All libraries received at least 1,207,605 read pairs passing filter (mean=2,766,149 read pairs, sd=1,582,161 read pairs). Average base quality phred scores for samples ranged from 33.6 to 34.8 (mean=34.4, sd=0.29). An average of 93% of all read pairs both passed the Trimmomatic filter, whereas 5.2% of all read pairs had either the forward or reverse read removed, and 1.8% had both members removed. Because our insert size was mostly larger than 300bp (which is two times the read length), fastq-join did not merge most reads—percentages of joined reads ranged from 24.0% to 35.1% for the different samples. Nuclear sequence divergence between the Mexican axolotl (the species from which probes were designed) and California tiger salamander in the exon targets averaged 1.84%.

### Reference assembly and read mapping

A total of 78,674,304 reads (all of the reads from the CTS individual) representing 11,960,279,114 bp were supplied to ARC for *de novo* assembly of targets. An average of 905 reads in iteration 1, 1,496 reads in iteration 2, 1,999 reads in iteration 3, 4,485 reads in iteration 4, 8,132 reads in iteration 5, and 11,199 reads in iteration 6 were assigned to each target for *de novo* assembly. The final assembly, after six iterations of the ARC assembly pipeline, contained 120,617 sequences for a total of 69,873,191 bp. After blasting the target sequences to the assembly and *vice versa*, we found a total of 8,386 RBBHs, or 96.3% of all targets. These assembled target contigs were 1,409 bp on average, for a total reference length of 11,813,341 bp. This average extension of 1,119 to each target sequence was expected, as the insert size in our genomic library preparations ranged up to roughly 550 bp. Thus 550 bp fragments that contained target sequence on either end could still be hybridized by the capture probes and their sequence at the other end recruited into the target assembly. Self-blasting the target RBBHs to one another resulted in 1,060 targets that also had hits with other targets. A total of 361,949 bp of such overlap was found between targets, and the overlapping bases were replaced with N’s to reduce the effects of repetitive sequences and chimeric assemblies.

An average of 18.21% of all reads across samples mapped to the chimera-masked reciprocal blast hit target assembly. Individual sample raw read mapping rates ranged from 6.7% to 34.8% (Table 2). The percentage of PCR duplicates present also varied widely across samples, ranging from 8.5% to 48.6% (mean=24.5%, sd=11.7%). After subtracting PCR duplicates from mapped reads, the percentage of unique reads on target varied between 5.4% and 30.8%, with a mean of 14.0% and standard deviation of 4.4% (Table 2).

Target-level metrics indicated that some targets performed significantly better than others (Figure 5). To control for variation in the number of reads received between samples, all libraries had their depths corrected to what would have been observed if they had received the same number of reads as the least-sequenced library in this study. To give an idea of the sequencing effort required to generate the depths listed below, this corresponded to approximately 2.4 million 150 bp reads against just over 2.5 million bp of total target sequence. Among all libraries, the average depth across target sequences was 7.99 (sd=3.33), and the average for the highest 100bp window within targets was 9.50 (sd=3.89). A total of 5,648 targets had a sequencing-effort corrected average depth across the target region greater than 5, and 2,283 had average depths greater than 10. For the 100 bp windows with the greatest depth for each target, 6,100 had depths greater than 5 and 3,313 had depths greater than 10.

**Figure 5.**
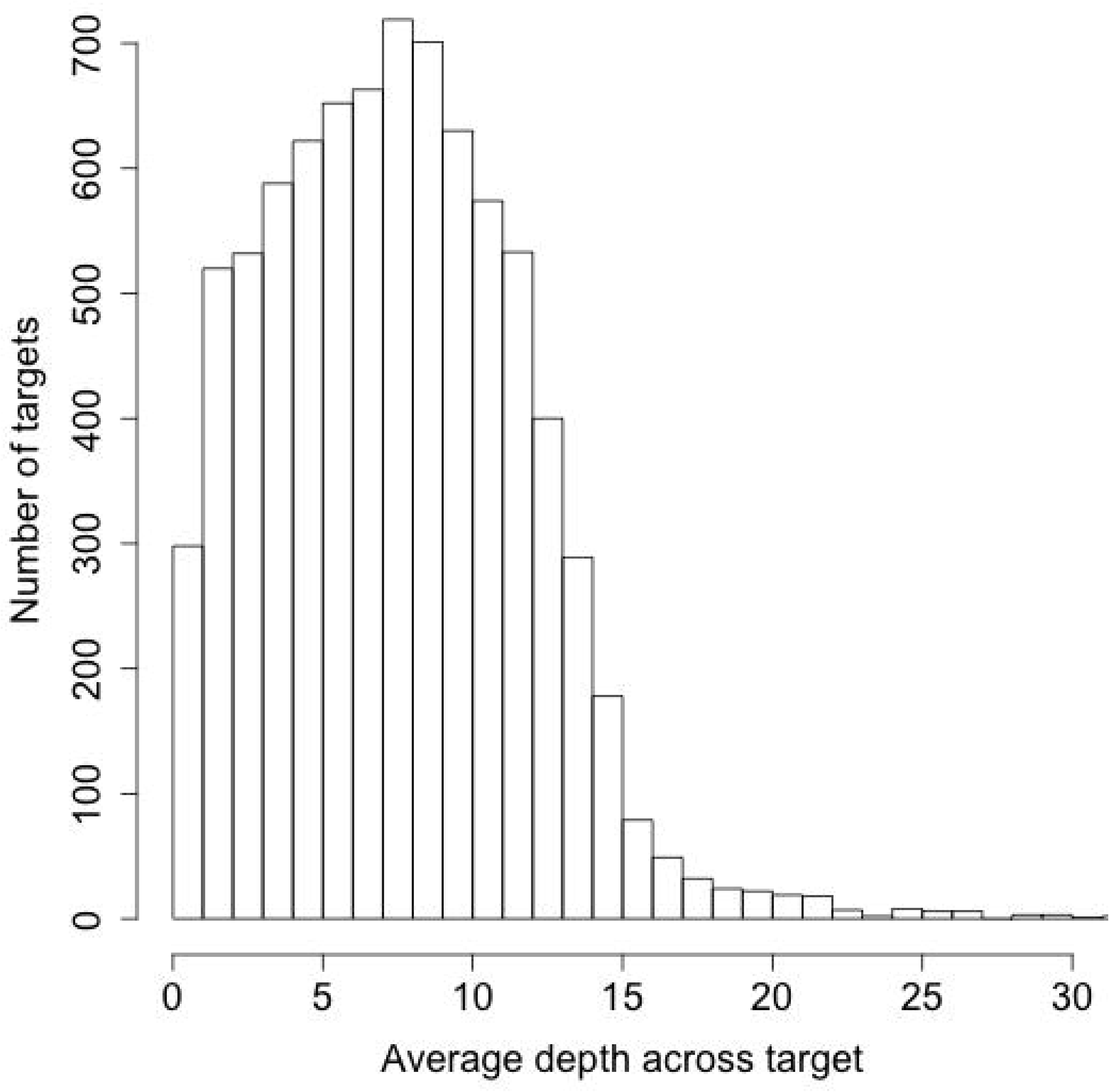
Average sequencing depths across targets. The average sequencing depth across all targets regions averaged between all samples, calculated using *samtools depth.* The highest 31 values, which had depths higher than 30, are not shown here.

### Effects of c_0_t-1 and input DNA amount in capture reactions

All models that incorporated the identity of the individual DNA pool underperformed (higher AIC value) nested models that did not incorporate information regarding the identity of the input DNA. Because of this, and because slope coefficients for the identity term in all models was never significant (p = 0.44 or greater), the identity of the individual did not significantly impact capture efficiency or read mapping, and models including this variable are not included in the summary tables.

Increasing the amount of individual input DNA and the amount of c_0_t-1 blocker were both associated with higher percentages of unique reads on target and higher realized sequence depth across targets (Tables 3 and 4, Figure 6). Linear regression recovered positive and significant slopes for both variables separately and when combined in multiple linear regression. Models predict an extra 1% unique reads on target for every 166 ng of extra individual input DNA (p = 0.000672) or every 6,750 ng of extra c_0_t-1 blocker (p = 0.00896) used in enrichment reactions. Regression coefficients for models that contained both individual input DNA and c_0_t-1 were quite similar to the single-variable model, differing by less than 3%. Individual input DNA and c_0_t-1 did a better job predicting the percentage of unique reads on target than the average depth across target regions (adjusted R^2^ of 0.325 vs 0.252 for the combined models). Finally, the models that contained both input DNA and c_0_t-1 as variables had better AIC scores and R^2^ values than the nested single-variable models (see Figure 7), and within the single-variable tests individual input DNA models outperformed c_0_t-1 models for both success measures in AIC and R^2^ (Tables 3 and 4).

**Table 3.** Model comparison predicting percentage of unique reads on target, sorted by AIC values *** signifies p < 0.001, ** signifies 0.001 < p < 0.01, * signifies 0.01 < p < 0.05.

| Model | $R^2$ | Adj. $R^2$ | AIC |
| --- | --- | --- | --- |
| $c_0t1^{**} + \text{inputDNA}^{***}$ | 0.3252 | 0.2982 | -193.6057 |
| $\text{inputDNA}^{***}$ | 0.2046 | 0.189 | -186.8963 |
| $c_0t1^{**}$ | 0.1265 | 0.1094 | -181.9297 |

**Table 4.**
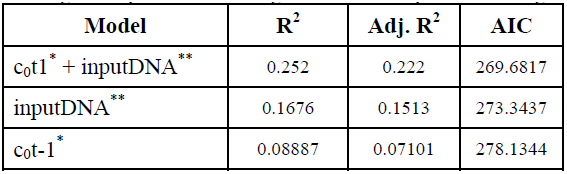
Model comparison predicting average depth across target region, sorted by AIC values *** signifies p < 0.001, ** signifies 0.001 < p < 0.01, * signifies 0.01 < p < 0.05.

**Figure 6.**
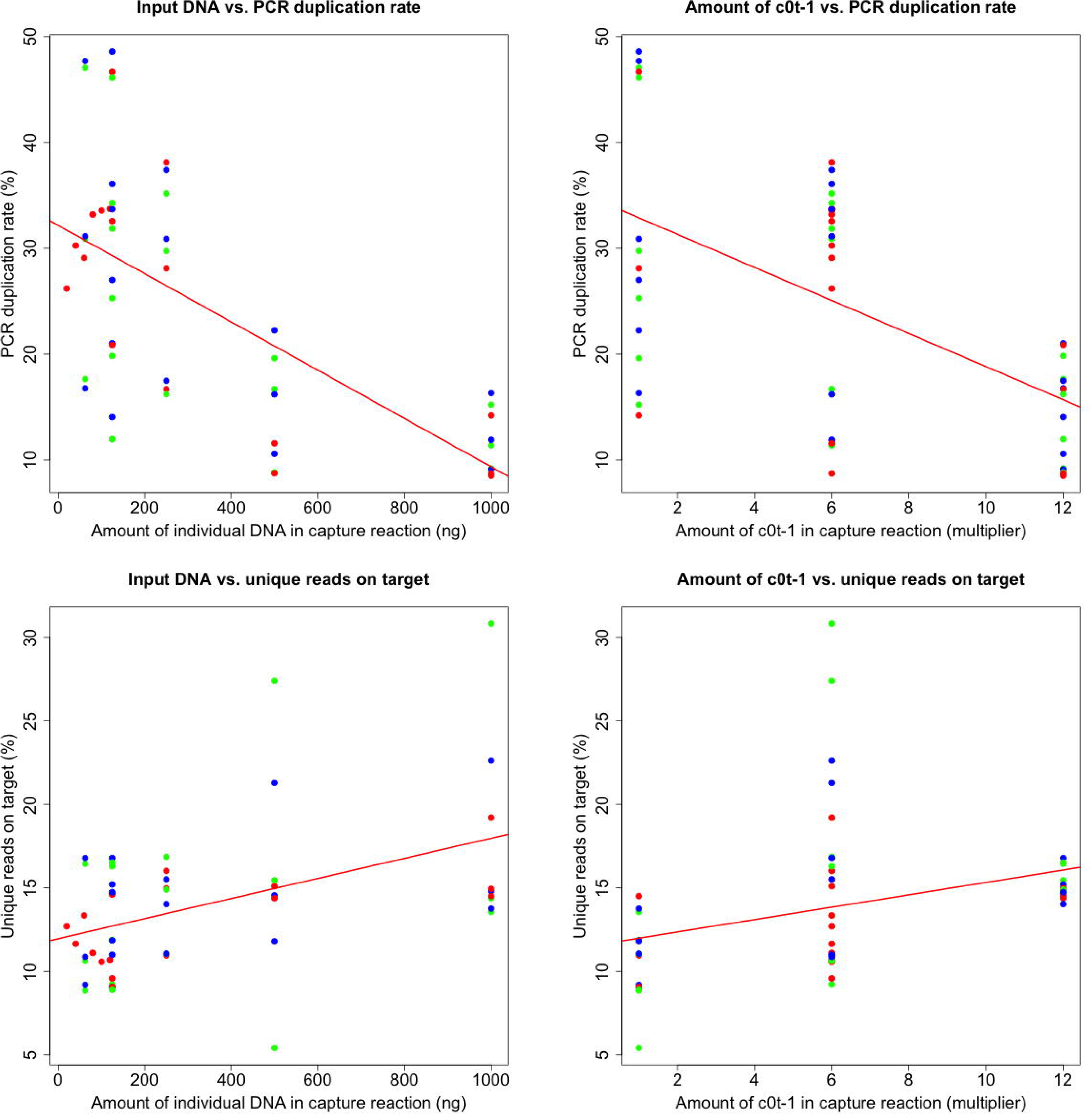
Relationship between individual input DNA and c_0_t-1 amounts to PCR duplication rates and percentages of unique reads on target. Each dot is an individual library: blue=CTS, green=F1, red=BTS*. P-values for slope coefficients in the four panels are: top left p = 1.39x10^−7^, top right p = 9.28x10^−6^, bottom left p= 0.000672, bottom right p = 0.00896.

**Figure 7.**
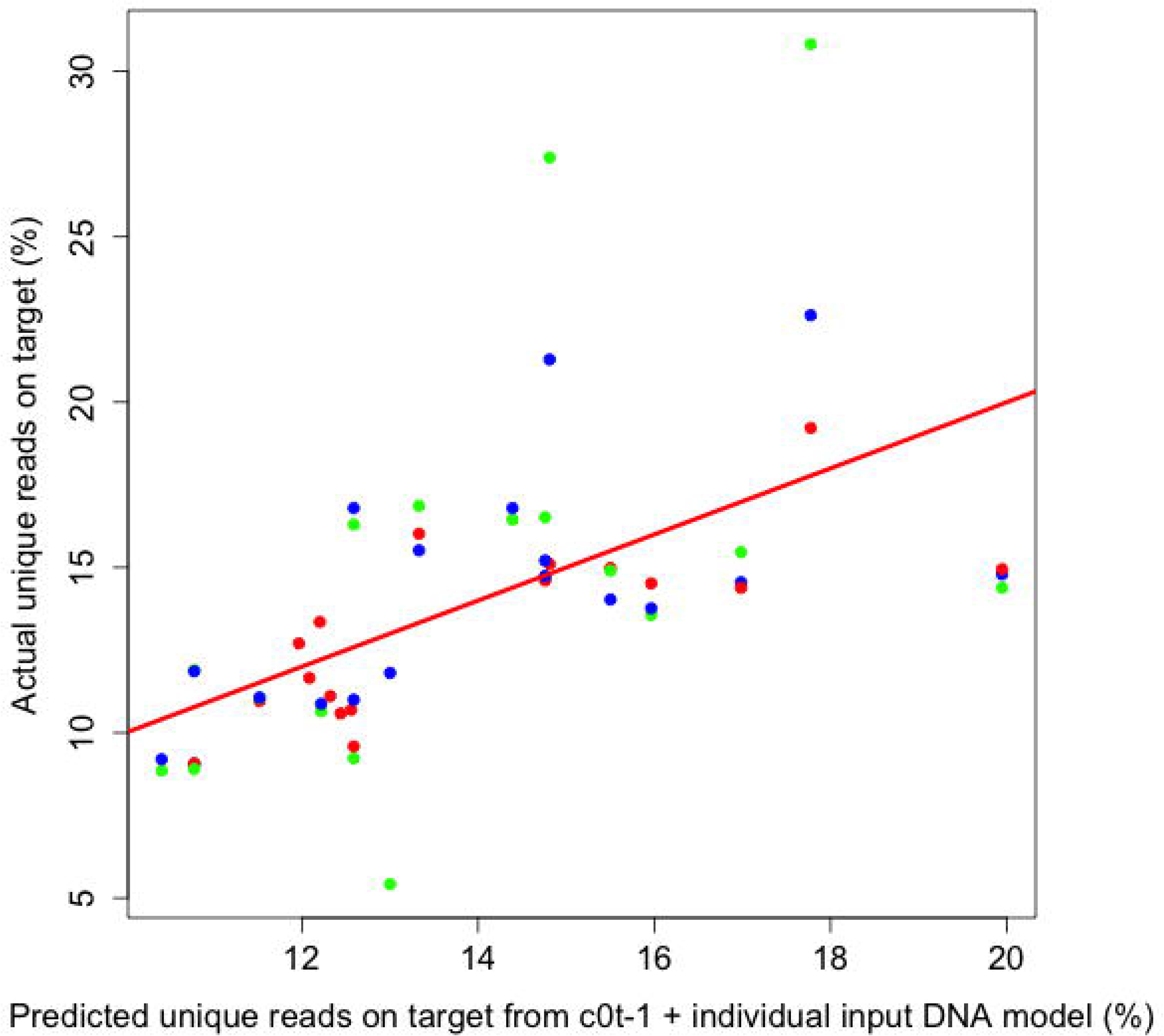
Predicted vs. actual unique reads on target using two-variable model. The model contains both c_0_t-1 and individual input DNA. Points close to the line mean that their unique reads on target are well-predicted by the two variables, and points farther away from the line are not as well predicted. Each dot is an individual library: blue=CTS, green=F1, red=BTS*.

## Discussion

Perhaps the most important conclusion from this experiment is that target capture experiments can indeed be successful in large-genome amphibians. This was not at all obvious based on prior work on these organisms, and our hope is that others will use these results to bring amphibians into the realm of population and phylogenomic analyses. The percentage of unique reads on target is the most important summary metric for enrichment, as it is essentially one minus the high quality data from the sequencer that is discarded. Our average percentage of unique reads on target across all library treatments was 14%; only three libraries were under 9%, while our four best-performing libraries were all over 20%. These numbers suggest that it is reasonable to sequence 50 to 100 samples on a single HiSeq lane for a capture array size similar to ours (2.5 megabases), depending on array configuration and coverage requirements.

Our rates of unique reads on target are in line with several other non-model exon capture studies for species with smaller genomes. For instance, Hedtke *et al.* designed Agilent probes from the *Xenopus tropicalus* genome and enriched libraries from two smaller-genome frogs, achieving rates of 7.4% unique reads on target in *Pipa pipa* and 47.8% in *Xenopus tropicalus* [9]. Bi *et al.* recovered 25.6% to 29.1% unique reads on target for an exon capture study in chipmunks [7]. Similarly, Cosart *et al.* designed an Agilent exon capture microarray from the *Bos taurus* genome and attained 20%-29% unique read mapping percentages in *Bos taurus*, *Bos indicus*, and *Bison bison* for a similarly-sized target array as this study [35]. Finally, Neves *et al.* reached 50% raw mapping rates in multiplexed exon capture experiments in *Pinus taeda*, a pine species with a roughly 21 gb genome (approximately 2/3 of the size of the salamander genomes in this study), although they did not report percentages of unique reads on target or levels of PCR duplication [8]. Several factors may be important in explaining these results, including a potential negative relationship between the phylogenetic distance to the species from which the capture array was developed and the percentage of unique reads on target, and the size of the genome under investigation. As more target capture studies are reported across diverse non-model taxa, we will better understand the relationship between genome size and enrichment efficiency, as well as the effects of designing capture probes from divergent taxa.

Human exome capture studies, which typically use predesigned sequence capture arrays across one of several different technologies (e.g. Truseq, Nimblegen, Agilent, or Nextera exome capture kits) often attain percentages of unique reads on target in the range of 40% to 70% or higher [36, 37]. This suggests that working from a well-assembled genome of the study species helps to increase the number of reads on target substantially. However, the high numbers in human experiments are likely also a function of the technologies used and the many iterations of probe set optimization experiments that have been conducted, and these may not be feasible in non-human systems.

We found evidence that increasing c_0_t-1 and individual input DNA into sequence capture reactions increased the percentage of unique reads mapping to targets in large-genome salamanders. As can be seen in Figure 6, this effect was driven largely by the correlation of these two variables with the reduction in PCR duplication rates. Because duplicate reads (reads with the same 5’ and 3’ mapping coordinates) are typically removed prior to genotyping analyses, lowering duplication rates as much as possible is critical for increasing the efficiency, and therefore reducing the sequencing costs of target enrichment studies. In addition to considering the variables tested here, researchers should also consider paired-end sequencing whenever possible in exon capture studies, as single-end reads have a much higher false identification rate of PCR duplication [38].

The low yields of DNA after enrichment and PCR are interesting. We speculate that they may be a consequence of libraries prepared from large genomes containing relatively low absolute numbers of on-target fragments in the pools during enrichment, so that a higher percentage of the pool is washed away. While qPCR of pre- and post-enrichment libraries using primers meant to amplify targeted regions is a useful way to test enrichment efficiency, we found that post-enrichment DNA concentrations may also be informative as to whether or not enrichment was successful for large-genome amphibians with this protocol (Figure 4). Also, we note that Library #18, which had a very high post-enrichment post-PCR DNA concentration, showed correspondingly low performance in terms of percentage of raw and unique reads on target (5.4% unique read mapping rate). This suggests that for this reaction, off-target fragments may not have been efficiently removed during the post-enrichment washing steps.

After duplicate removal, we observed a greater than five-fold difference in unique read mapping percentages (from 5.4% to 30.8%) among the samples tested in this experiment. While even the low end of our enrichment efficiency values are encouraging for future exon capture studies in large-genome amphibians, regularly attaining unique reads on target percentages at the upper end of our success rate would lead to a concurrent 5X reduction in sequencing costs for a given target coverage depth. In the future, we would like to test the effects of increasing the amount of DNA (and therefore number of genome copies) used for library preparations, as well as increasing the total amount of DNA in a single enrichment reaction above the 1,000 ng used here, with the hope that both of these steps will further reduce PCR duplication rates.

## Conclusions

Exon capture is a viable technology for gathering data from thousands of nuclear loci in large numbers of individuals for salamanders and other taxa with large (at least 30 gb), highly repetitive genomes. We recommend using at least 30,000 ng of species-specific c_0_t-1 blocker, and as much input DNA as possible for each individual multiplexed into a capture reaction when working with large-genome species. Ongoing research in our lab to further optimize large genome target capture is focusing on the tradeoffs of different multiplexing regimes and the tradeoffs from increasing the total amoung of DNA going into capture reactions and individual library preparations (Figure 1). Although we can only speak directly to experiments that utilize custom MYbaits exon enrichment reactions, we see no reason why our results should not generalize to other platforms such as UCEs [2].

As large-scale sequencing projects become the norm for data acquisition in non-model systems, it is crucial to build a body of literature with standard reporting metrics for both laboratory procedures and data filtering and analysis. Gathering information about best practices in custom array target enrichment from experiments in the literature is difficult due to the lack of standardization in reporting metrics. At a minimum, we suggest that researchers report raw mapping rates to target sequences, PCR duplication rates (ideally based on paired-end reads), and average depths across the different targets, including standard deviations, for a given sequencing effort. Standardized metrics will allow researchers to evaluate whether a particular probe set may work in their study system and how much sequencing may be needed. We hope that this study can help set a precedent for such reporting on successful laboratory procedures, including a thorough discussion of efficiency and success of target capture in non model organisms.

## Availability of supporting data

The data set supporting the results of this article is available at Genbank:PRJNA285335. The target sequences used for this study, the corresponding *Ambystoma mexicanum*-derived capture probes, and the source code used to analyze the data from this experiment are available at http://dx.doi.org/10.5281/zenodo.18587 [39].

## Competing interests

The authors declare that they have no competing interests.

## Authors′ contributions

EMM contributed to the design of the study, performed some of the molecular work, analysed the results, and wrote the manuscript. GGM contributed to the design of the study, performed most of the laboratory work, and revised the manuscript. HBS contributed to the design of the study and interpretation of results, and revised the manuscript. All authors read and approved the final manuscript.

## Acknowledgements

We thank Randal Voss for the *Ambystoma mexicanum* sequences used to design the capture array, and Brant Faircloth for input on experimental design and laboratory troubleshooting. Animal work was conducted under California Department of Fish and Wildlife permit #SC- 2480 and associated MOU, USFWS permit #TE-094642-9, and UCLA IACUC protocol #2013-011. This experiment used the Vincent J. Coates Genomics Sequencing Laboratory at UC Berkeley, supported by NIH S10 Instrumentation Grants S10RR029668 and S10RR027303. EMM and HBS are supported by NSF-DEB 1257648.

